# Diet context gates AgRP neuron involvement in Semaglutide-induced weight loss

**DOI:** 10.1101/2025.07.13.664597

**Authors:** Mateus d’Ávila, Roberto Collado-Pérez, Zhongwu Liu, Tamas L. Horvath

## Abstract

Semaglutide, a GLP-1R agonist, is widely used for obesity and type 2 diabetes, but its neural mechanisms remain unclear. AgRP neurons regulate energy balance, yet their role in Semaglutide’s effects is unknown. We show that sustained treatment of female mice with Semaglutide leads to activation rather than inhibition of AgRP neurons. Ablation or hypofunction of AgRP neurons through cell-specific knockout of *Sirt1* reduces Semaglutide-induced weight loss and impairs its hypoglycemic effects in female mice under Standard Diet. However, acute or chronic exposure to High-Fat Diet makes AgRP neurons dispensable for weight loss, suggesting that neural substrates for the actions of Semaglutide depends on dietary composition. Re-exposure to Standard Diet recovers the necessity for AgRP neurons, underscoring the influence of nutritional status on GLP-1R pathways. Our findings show the necessity for AgRP neurons in sustaining Semaglutide-induced weight loss in female mice on standard diet *in vivo*.

## Main

Obesity is a major global health concern, associated with increased risk of metabolic disorders, cardiovascular disease, and mortality^1^. Despite its complex etiology, pharmacological interventions have emerged as effective strategies for weight management^2,3^. Among these, Glucagon-like peptide-1 receptor agonists (GLP-1RA), such as Semaglutide, have demonstrated potent hypoglycemic and weight-loss effects, making them a cornerstone in obesity treatment^4,5^. However, the precise mechanisms by which GLP-1R agonists sustain long-term weight loss remain incompletely understood, limiting the ability to optimize their therapeutic efficacy.

Agouti-related peptide (AgRP) neurons, located in the Arcuate Nucleus of the hypothalamus, are key regulators of energy homeostasis, integrating peripheral metabolic signals to drive feeding behavior^6-9^. While GLP-1R agonists are known to promote weight loss, whether these effects require AgRP neuron activity remains unresolved. The literature presents conflicting findings, with some studies suggesting that AgRP signaling plays a role in the effects of GLP-1R agonists^10-13^, while others argue that GLP-1RAs do not act on AgRP neurons and that the arcuate nucleus is dispensable for their metabolic effects^6,14,15^. Despite these conflicting reports, no study has directly assessed the functional role of AgRP neurons in mediating Semaglutide’s effects *in vivo*. Given that diet composition itself can influence the neural circuits regulating energy balance - with high-fat diets engaging mesolimbic reward pathways while lower-fat, fiber-rich diets primarily recruit hypothalamic homeostatic circuits - it is possible that the involvement of AgRP neurons in Semaglutide’s effects depends on dietary context^16,17^. Understanding whether AgRP neurons play a necessary role in Semaglutide-induced weight loss and metabolic adaptation remains a critical gap in the field.

To investigate this, we employed a cell-specific deletion of *Sirt1* in AgRP neurons, a model previously shown to impair AgRP activation without altering baseline activity. Additionally, we used a transgenic model expressing diphtheria toxin receptor (DTR) in AgRP neurons, allowing for their selective ablation to further confirm our findings. We investigated the effects of Semaglutide on AgRP neuronal activity in different time points of treatment, and assessed the participation of adrenergic β3 receptor signaling to its sustained weight loss effects. We then examined the role of AgRP neurons in mediating the effects of Semaglutide under standard and high-fat diet conditions, in both lean and diet-induced obese mice, to determine whether dietary composition or metabolic state influences AgRP neuron dependence. By integrating molecular, pharmacological, electrophysiological and dietary manipulations, our study aims to provide deeper insight into how AgRP neurons contribute to GLP-1R agonist-mediated weight loss and whether their role is modulated by diet-induced shifts in feeding circuitry.

## Results

### AgRP neurons are necessary for Semaglutide-induced weight-loss in female mice under Standard Diet through an adrenergic β3 receptor-dependent mechanism

Initially, we assessed the effects of semaglutide on lean AgRP-*Sirt1* wild-type (WT) and knockout (KO) mice under Standard Diet (SD) (Fig. 1A). We found that semaglutide induced significant acute weight loss in WT (*Cre*-negative) mice, while this effect was attenuated in KO animals (Fig. 1B). Sustained treatment led to a complete rebound of body weight to baseline levels in KO but not in WT (Fig. 1B). Analysis of body composition through fMRI revealed that WT and KO animals significantly lose fat mass after 2 days of treatment with Semaglutide in comparison to their correspondent vehicle controls (Fig. 1B). However, sustained treatment for 15 days leads to increase in fat mass levels in KO but not in WT mice (Fig. 1B), supporting the necessity for functional AgRP circuitry to sustain fat loss. These effects were sex-specific, as male KO mice exhibited similar weight loss to WT ones (Supplementary Fig. 1A), leading us to proceed with further experiments in females only. To further support our findings, we assessed body weight in AgRP^DTR^ animals treated with Semaglutide and observed a similar pattern of weight regain as the one observed in AgRP-Sirt1 KO animals (Supplementary Fig. 1B). Our data also shows that semaglutide reduced caloric intake in both WT and KO mice, but KO animals exhibited enhanced anorexigenic effect in comparison to their baseline (Fig. 1C). Despite differences in calorie intake, levels of energy expenditure (Fig. 1D) and energy balance (Fig. 1E) decreased similarly in both groups relative to baseline, suggesting that AgRP circuitry is not involved in the effects of Semaglutide over expenditure.

**Fig. 1.**
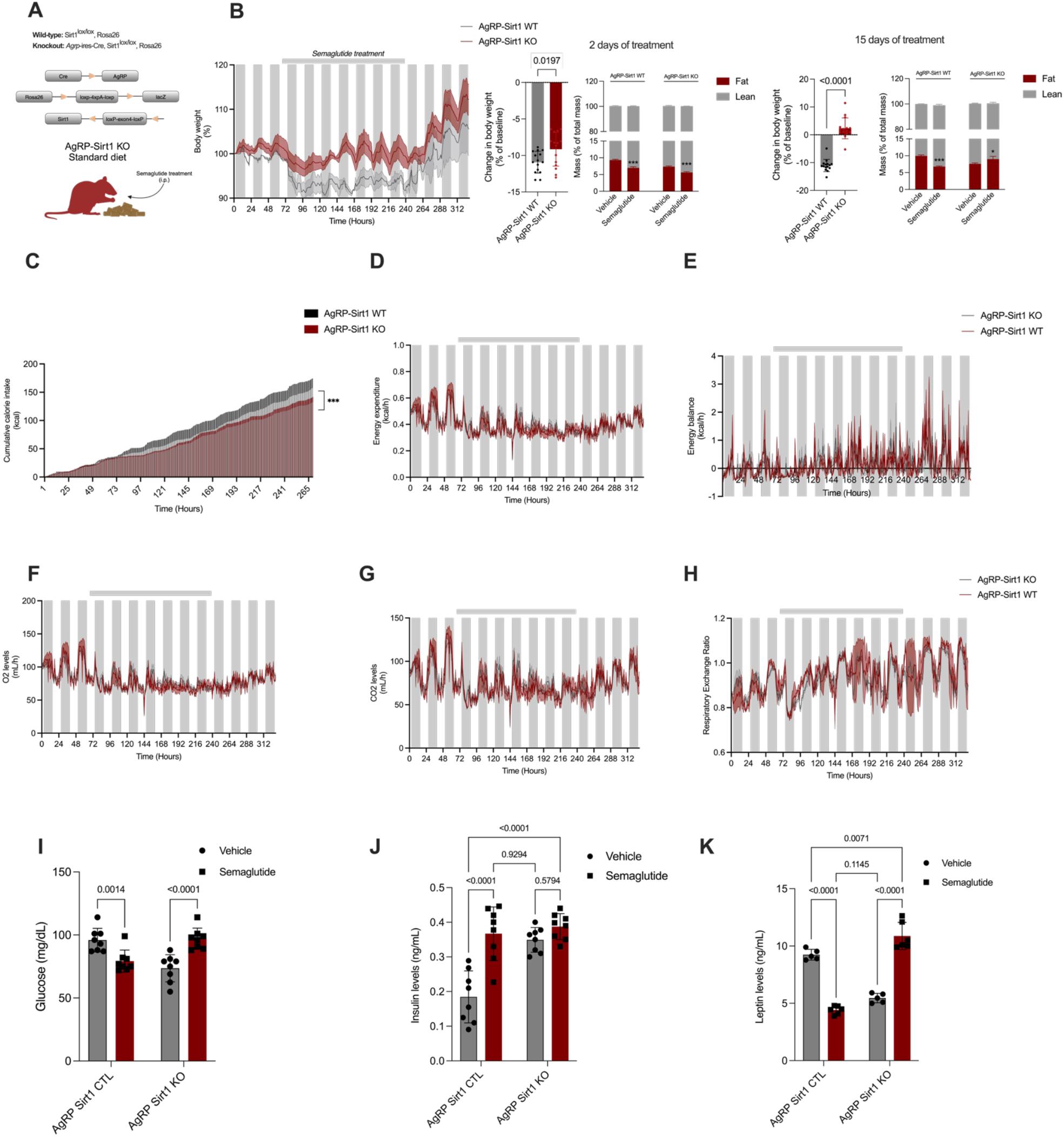
Female mice under Standard Diet require AgRP neurons for sustained weight loss and hypoglycemic effects. **(A)** Schematic representation of the *Agrp*-ires-*Cre*-*Sirt1*^*lox/lox*^ mice used in the study. WT mice lack the expression of *Agrp*-ires-*Cre*, **(B)** Left, percent change in body weight between Control and AgRP-Sirt1 Knockout (KO) mice in response to treatment with Semaglutide. Right, percent of change in body weight and body composition of mice treated with semaglutide for 2 or 15 days, **(C)** Cumulative food intake of WT and KO animals treated with Semaglutide, **(D)** Energy expenditure (kcal/h) of WT and KO animals, **(E)** Energy balance of WT and KO animals before, during, and after treatment with Semaglutide, **(F)** O_2_ levels of WT and KO animals, **(G)** CO_2_ levels of WT and KO animals, **(H)** Respiratory exchange ratio of WT and KO animals, suggesting changes in substrate utilization; **(I)** Glucose, **(J)** Insulin, and **(K)** Leptin levels of WT and KO mice treated with vehicle or with Semaglutide for 15 days, Silver bars represent the period in which mice received Semaglutide treatment. Details on statistical analysis can be found at supplementary Table 1. *p<0.05 vs AgRP-*Sirt1* KO Vehicle; ***p<0.001 vs AgRP-*Sirt1* WT Vehicle.

To further explore metabolic adaptations, we assessed O_2_ and CO_2_ levels, and analyzed respiratory exchange ratio (RER) as a proxy for substrate utilization, as reported elsewhere (18). Following the first Semaglutide injection, both WT and KO mice displayed a steep reduction in O_2_ and CO_2_ levels (Fig.s 1F and 1G, respectively) and in RER in comparison to baseline levels, indicating a shift towards fat oxidation (Fig. 1H). However, with prolonged treatment, both groups gradually returned to an RER of ∼1.0 during the dark phase (Fig. 1H) while remained with low O_2_ and CO_2_ levels relative to baseline, implying a maintenance of carbohydrate utilization under a state of hypometabolism.

**Table 1.**
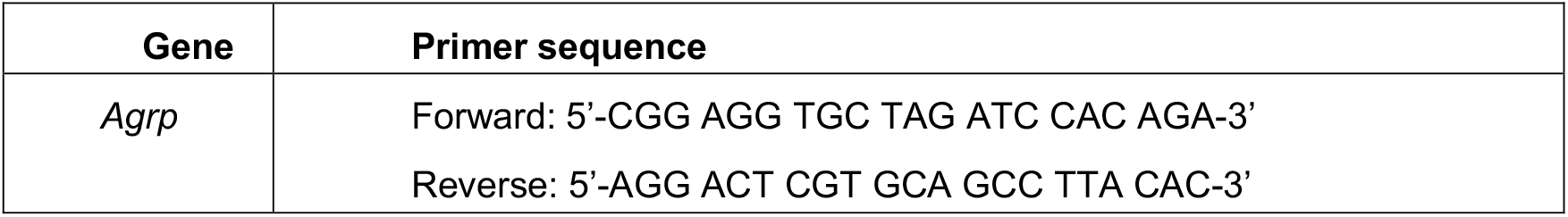
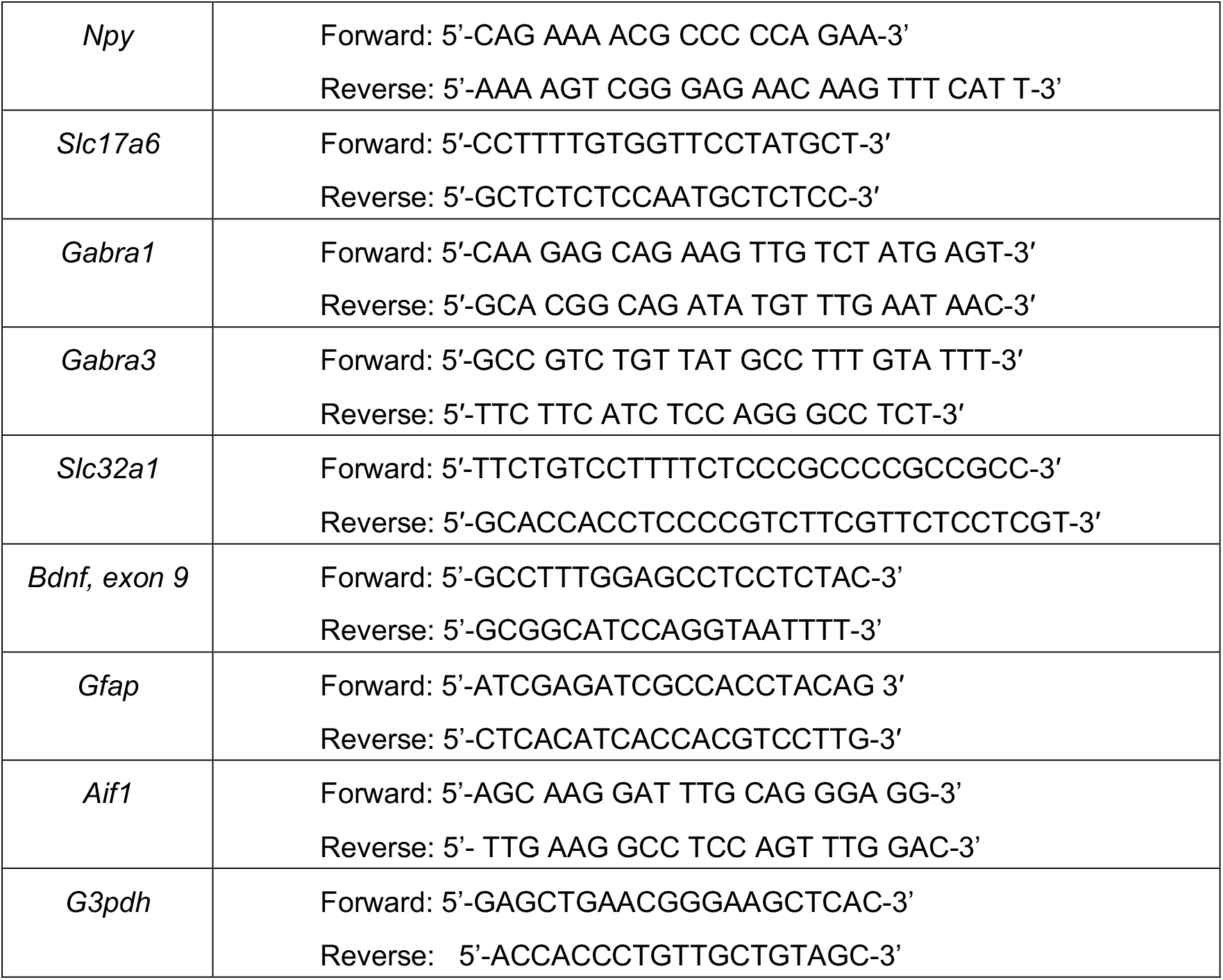
Primers used for RT-qPCR

Our data also shows that sustained Semaglutide treatment lowered blood glucose levels in WT mice but failed to do so in KO animals (Fig. 1I), supporting the necessity of AgRP neurons for the hypoglycemic effects of GLP-1RA in lean mice. AgRP^DTR^ animals also displayed impaired hypoglycemic response to Semaglutide, further corroborating the necessity of these cells for sustained low glucose levels (Supplementary Fig. 1C). Alterations in glucose profile of AgRP-*Sirt1* mice were accompanied by an increase in circulating insulin in WT’s but not in KO’s (Fig. 1J). Notably, KO mice displayed higher baseline insulin levels, which may reflect underlying alterations in glucose homeostasis. One possible explanation for the lack of further insulin increase in KO mice is that their pancreatic β-cells might already be secreting insulin at near-maximal capacity, limiting the insulinotropic effects of Semaglutide in this model. Additionally, leptin levels decreased in WT mice following sustained Semaglutide treatment, whereas KO mice exhibited an increase in circulating leptin (Fig. 1K). Given that leptin levels typically correlate with adiposity, this finding is likely a reflection of the fat mass rebound observed in KO mice (Fig. 1B), which in turn might lead to elevated leptin secretion, further supporting the role of AgRP neurons in mediating Semaglutide’s long-term metabolic effects on adiposity.

Our data showing hypofunction of AgRP neurons mitigating Semaglutide’s weight-loss effect implies that sustained treatment results in activation rather than inhibition of AgRP cells. To further support this, gene expression analysis revealed no changes in *Agrp* mRNA levels after 2 days of Semaglutide treatment, while *Npy* expression was reduced (Fig. 2A). However, after 15 days, both *Agrp* and *Npy* were upregulated, suggesting a time-dependent activation of AgRP neurons (Fig. 2A).

**Fig. 2.**
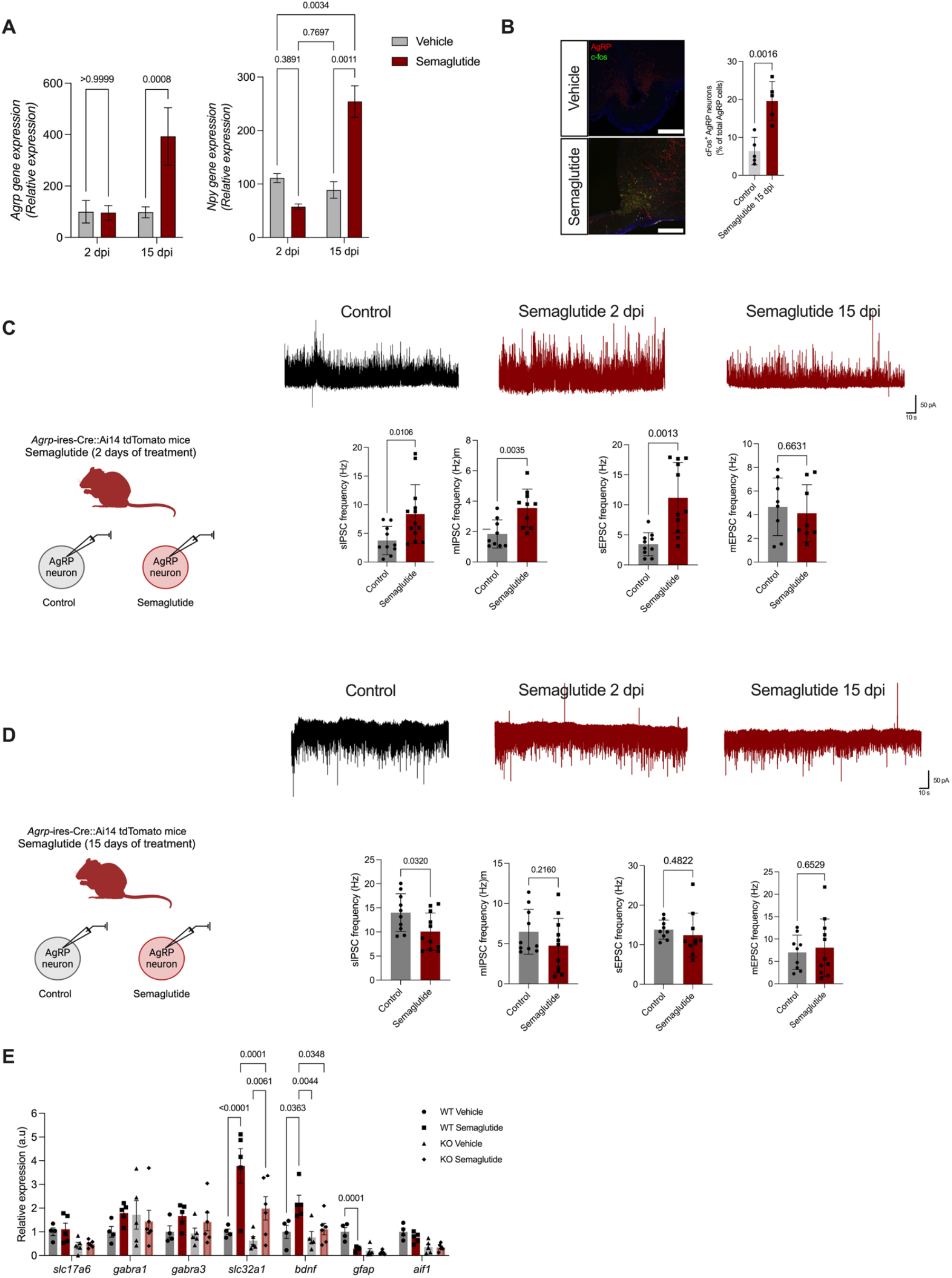
Sustained treatment with Semaglutide increases the activity of AgRP neurons. **(A)** *Agrp* and *Npy* relative gene expression of animals treated with vehicle or Semaglutide for 2 or 15 days (dpi = days post-injection), **(B)** Immunostaining showing c-fos (Green) and AgRP-tomato (Red) co-localization in the Arcuate Nucleus of mice treated with Semaglutide for 15 days; white bar: 50 μm **(C)** Left, schematic representation of whole-cell patch clamp recordings of AgRP-Ai14 *tdTomato* mice treated with Semaglutide for 2 days (Upper). Upper, representative sIPSC traces of control (Vehicle), Semaglutide 2 dpi and Semaglutide 15 dpi cells. Histograms show frequency (Hz) of spontaneous and miniature inhibitory and excitatory postsynaptic currents (sIPSC, mIPSC, sEPSC and mEPSC respectively) of mice treated with vehicle or Semaglutide for 2 days, **(D)** Left, schematic representation of whole-cell patch clamp recordings of AgRP-Ai14 *tdTomato* mice treated with Semaglutide for 15 days (Upper). Upper, representative sEPSC traces of control (Vehicle), Semaglutide 2 dpi and Semaglutide 15 dpi cells. Histograms show frequency (Hz) of spontaneous and miniature inhibitory and excitatory postsynaptic currents (sIPSC, mIPSC, sEPSC and mEPSC respectively) of mice treated with vehicle or Semaglutide for 15 days. **(E)** Hypothalamic gene expression of AgRP-*Sirt1* WT and KO mice treated with vehicle or Semaglutide for 15 days. Closed circles: AgRP-*Sirt1* WT Vehicle, Closed Square: AgRP-Sirt1 Semaglutide, Closed triangles: AgRP-*Sirt1* KO Vehicle, Closed Diamond: AgRP-*Sirt1* KO Semaglutide. Details on statistical analysis can be found at supplementary Table 1.

Immunofluorescence confocal analysis further confirmed this, as sustained Semaglutide treatment led to increased c-Fos co-localization with AgRP neurons labeled with tdTomato (Fig. 2B), indicating enhanced neuronal activity. To gain deeper insight into this effect, we performed whole-cell patch-clamp recordings in animals treated with Semaglutide. After 2 days of Semaglutide treatment, AgRP neurons displayed increased frequency of spontaneous and miniature inhibitory postsynaptic currents (sIPSC and mIPSC, respectively) alongside an increase in spontaneous excitatory currents (sEPSC), suggesting a balanced increase in excitatory drive and inhibitory restraint (Fig. 2C). However, after 15 days, sIPSC was reduced while excitatory currents remained unchanged, indicating a shift toward net excitation (Fig. 2D). In line with this synaptic remodeling, Semaglutide increased hypothalamic expression of *Bdnf* and *Slc32a1* (Fig. 2E), markers of neural plasticity and GABAergic signaling, respectively, while reducing *Gfap* levels (Fig. 2E), suggesting enhanced plasticity, inhibitory tone, and reduced astrocyte reactivity. These transcriptional changes were absent in AgRP-*Sirt1* KO mice, indicating that AgRP circuitry is required for Semaglutide-induced neuroplasticity. We also tested whether hormonal alterations could be contributing to AgRP activation promoted by semaglutide. Semaglutide did not affect circulating ghrelin levels (Fig. S1D) and co-treatment with leptin enhanced rather than inhibited weight loss and decrease in fat mass (Fig. S1E and 1F, respectively), suggesting that AgRP activation is not being driven by those compensatory endocrine factors. In summary, our data shows that sustained treatment with Semaglutide leads to synaptic reorganization and activation of AgRP neurons, while impairment of their activity mitigates weight-loss effects.

Given the described role of AgRP neurons in the modulation of sympathetic signaling^18^, we co-treated mice with SR 59230A, an antagonist of β3-adrenergic receptor (β3-AR), and assessed their metabolic parameters. As predicted, blockade of β3-AR inhibited weight loss promoted by 7 days, but not 2 days (Data not shown), of administration with semaglutide (Fig. S2A), while increasing calorie intake (Fig. S2B) and energy expenditure (Fig. S2C). Assessment of O_2_ (Fig. S2D) and CO_2_ levels (Fig. S2D) shows that SR 59230A also inhibits the decrease promoted by semaglutide, as well as recovers baseline RER levels. Analysis of whole-body substrate utilization shows that Semaglutide increases fat metabolism (Fig. S2G) and decreases carbohydrate utilization (Fig. S2H), while co-treatment with β3-AR antagonist partially suppresses this effect, supporting that β3-AR is necessary for fat utilization evoked by Semaglutide (Fig. S2E).

### AgRP neurons are dispensable for Semaglutide-induced weight-loss in diet-induced obese mice or lean animals acutely exposed to High-Fat Diet

Given the widespread use of semaglutide for obesity management, and to further characterize the mechanism of action of this drug, we assessed the requirement of AgRP neurons to weight loss of diet-induced obese mice (Fig. 3A). Obese female KO mice displayed a similar weight loss level in comparison to WT animals treated with semaglutide (Fig. 3B), as well as did not show significant differences in fat loss (Fig. 3C), in contrast to what we observed in lean female mice. Male obese KO animals also displayed no major difference regarding weight loss or overall metabolic parameters in comparison to WT’s (Fig. S3).

**Fig. 3.**
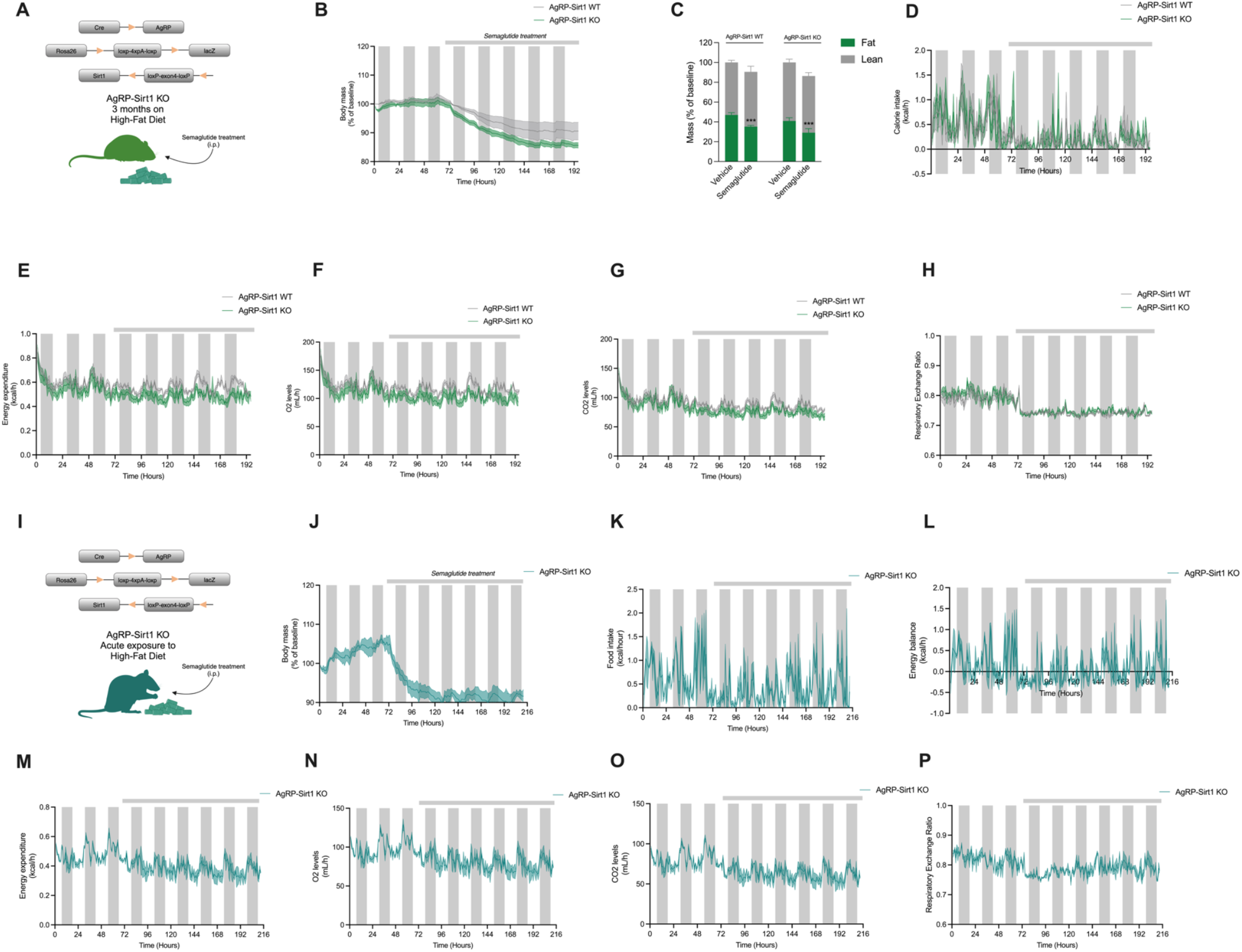
AgRP neurons are dispensable in obese mice or mice acutely exposed to High-Fat Diet. **(A)** Schematic representation of female obese mice used for this study. Animals were under 60% High-Fat Diet for at least 3 months before start of treatment with Semaglutide, **(B)** Percent change in body weight between WT and KO obese mice treated with Semaglutide, **(C)** Body composition of WT and KO mice treated with vehicle or with Semaglutide for 7 days, **(D)** Calorie intake (kcal/h) of WT and KO obese animals, **(E)** Energy expenditure (kcal/h) of obese WT and KO mice before and during Semaglutide treatment, **(F)** O_2_ and **(G)** CO_2_ levels of obese WT and KO animals treated with Semaglutide, **(H)** Respiratory exchange ratio of obese WT and KO animals before and during Semaglutide treatment. **(I)** Schematic representation of female KO mice acutely (3 days) exposed to 60% High-Fat Diet, **(J)** Percent change in body weight of KO animals before and during Semaglutide treatment, **(K)** Food intake (kcal/h) of KO animals before and during Semaglutide injections, **(L)** Energy balance (kcal/h) of KO animals before and during injections with Semaglutide, **(M)** Energy expenditure (kcal/h) of KO animals treated with Semaglutide, **(N)** O_2_ and **(O)** CO_2_ levels of KO animals acutely exposed to HFD treated with Semaglutide, **(P)** Respiratory exchange ratio of female mice acutely exposed to HFD and treated with Semaglutide. Silver bars represent the period in which mice received Semaglutide treatment. Details on statistical analysis can be found at supplementary Table 1. ***p<0.001 vs AgRP-*Sirt1* WT or KO Vehicle.

Semaglutide treatment reduced calorie intake in both WT and KO mice relative to baseline across five days of analysis (Fig. 3D). However, KO mice exhibited lower energy expenditure (Fig. 3E) and reduced O_2_ and CO_2_ levels compared to baseline (Fig. 3F and 3G, respectively). RER decreased similarly in both groups (Fig. 3H), indicating that Semaglutide promotes a shift to fat oxidation in obese mice through an AgRP-independent mechanism.

To determine whether the lack of AgRP circuitry involvement in Semaglutide-induced weight loss is driven by obesity-related physiological alterations or by diet exposure alone, we subjected lean female mice to HFD for three days before treatment (Fig. 3I). Strikingly, KO mice under short-term HFD exhibited no weight rebound (Fig. 3J), indicating that brief dietary fat exposure is sufficient to render AgRP neurons dispensable for Semaglutide’s effects. These animals also showed reduced calorie intake (Fig. 3K) and energy balance (Fig. 3L) relative to baseline. Semaglutide further decreased energy expenditure (Fig. 3M), O_2_ consumption (Fig. 3N), and CO_2_ production (Fig. 3O), while RER analysis confirmed a sustained shift to fat oxidation throughout the observation period (Fig. 3P).

### Reinstating Standard Diet reestablishes the requirement for AgRP neurons in sustaining Semaglutide-induced weight loss

To assess whether the dispensability of AgRP neurons under HFD could be reversed, we switched the diet of mice from HFD to SD (Fig. 4A), and monitored body weight along with other metabolic parameters. After one day of exposure to SD, animals started receiving Semaglutide treatment. Our findings show that KO but not WT animals show weight rebound after 4 days of Semaglutide injections (Fig. 4B), suggesting that re-exposure of mice to Standard Diet reestablishes the requirement for AgRP neurons in sustaining GLP-1RA-induced weight loss. As observed in lean mice under SD, switching the diet from high to standard fat levels leads to a transient suppression of calorie intake (Fig. 4C) and energy balance (Fig. 4D), which returns to baseline levels after 2 days of treatment in both WT and KO animals. Semaglutide treatment decreases energy expenditure in comparison to baseline independently of the genotype of mice (Fig. 4E), as well as O_2_, (Fig. 4F) and CO_2_ (Fig. 4G) levels. Analysis substrate utilization through RER shows that KO animals start metabolizing carbohydrates after 2 days of injection with Semaglutide (Fig. 4H), suggesting the necessity of these cells for reinstating carbohydrate utilization.

**Fig. 4.**
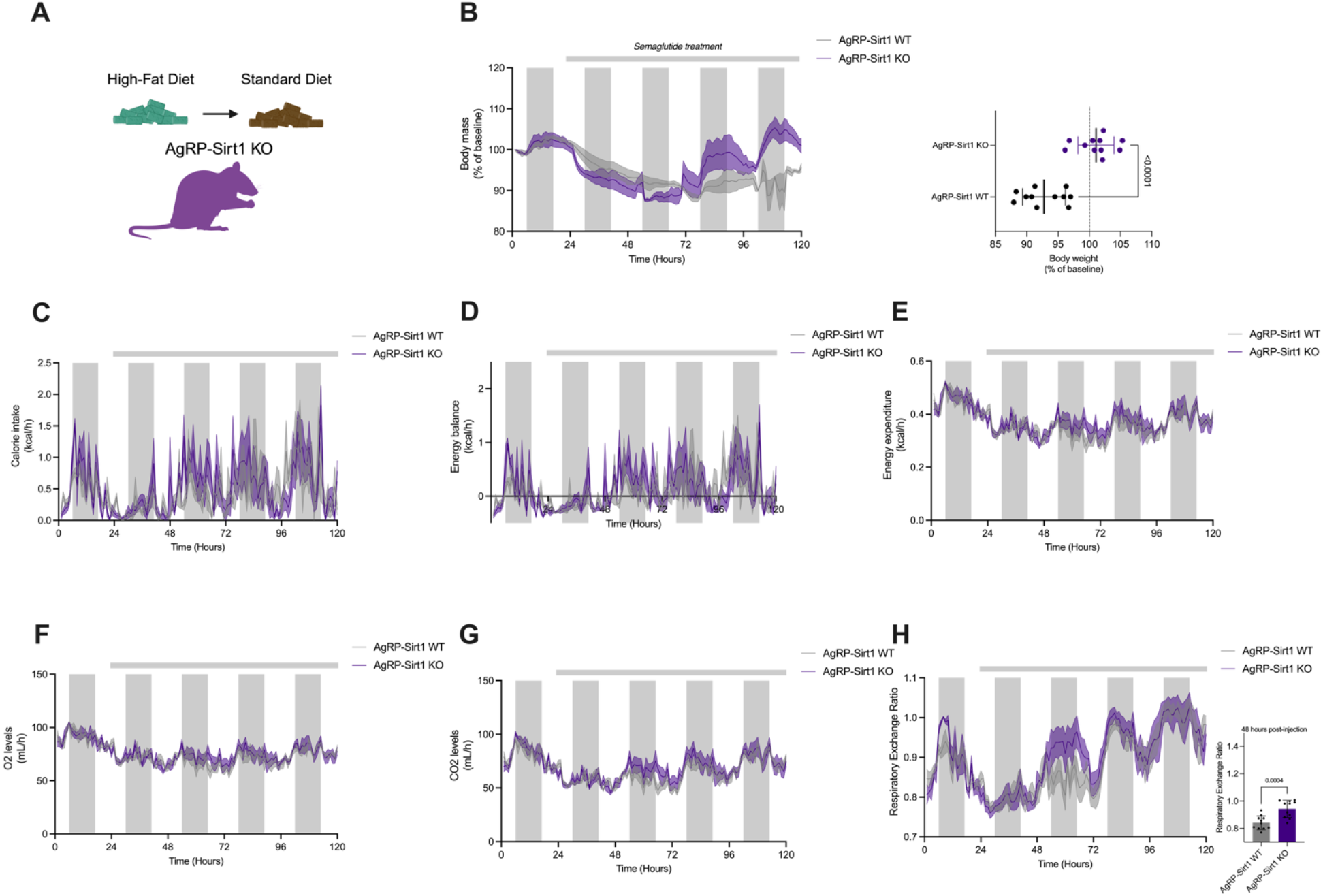
Re-exposure to Standard Diet reinstates AgRP neuron dependency for Semaglutide-induced weight loss. **(A)** Schematic representation of obese female mice who had their diet switched from High-to Low-fat and started receiving Semaglutide injections after 1 day, **(B)** Percent change in body weight of obese WT and KO animals re-exposed to Standard Diet and treated with Semaglutide, **(C)** Calorie intake (kcal/h) of WT and KO animals re-exposed to Standard Diet treated with Semaglutide, **(D)** Energy expenditure (kcal/h) of WT and KO animals re-exposed to Standard Diet treated with Semaglutide, **(E)** Energy balance (kcal/h) of WT and KO animals re-exposed to Standard Diet, **(F)** O_2_ and **(G)** CO_2_ levels of WT and KO animals re-exposed to Standard Diet and injected with Semaglutide, **(H)** Respiratory exchange ratio of WT and KO animals re-exposed to Standard Diet and treated with Semaglutide. Silver bars represent the period in which mice received Semaglutide treatment. Details on statistical analysis can be found at supplementary Table 1.

## Discussion

Our findings demonstrate that the involvement of AgRP neurons in Semaglutide-induced weight loss is modulated by dietary composition. Under standard chow conditions, AgRP neurons are required to mediate the GLP-1RA sustained effects on body weight in female mice. However, a shift to a high-fat diet disrupts this dependency, enabling Semaglutide to promote weight loss independently of AgRP circuitry. Reversing the diet from HFD to SD reengages AgRP neurons in weight-loss, strengthening the hypothesis that the neural pathways engaged by Semaglutide are dynamically shaped by the nutritional environment, highlighting the importance of diet in modulating the central mechanisms of GLP-1RA weight regulation.

Sustained treatment with Semaglutide paradoxically activates AgRP neurons, and this activation is essential for its weight loss and hypoglycemic effects. This challenge prior models on the mechanism of action of GLP-1RA describing an inhibitory effect over AgRP neurons as a drive for its anorexigenic and weight loss effects^10-12,19^. The discrepancy between our findings and previous reports may stem from differences in methodological approaches, as earlier studies employed acute *ex vivo* preparations to assess immediate actions of GLP-1RA on AgRP neurons, which fail to capture the dynamic, metabolic state-dependent plasticity of hypothalamic circuits^10,11,19^. Consequently, acute cellular inhibition observed in these conditions may reflect an immediate pharmacological effect of GLP-1RA rather than its integrated response to sustained energy imbalance. By assessing the electrophysiological properties of AgRP neurons from animals treated with Semaglutide, our work captured the synaptic adaptations in this circuit in response to sustained energy imbalance, particularly a reduction in inhibitory inputs to AgRP cells, suggesting a shift towards net excitation (Fig. 2D), which aligns with the observed increase in *Agrp* and *Npy* gene expression (Fig. 2A), as well as c-fos staining (Fig. 2B). These electrophysiological changes were accompanied by an AgRP-*Sirt1*-dependent increase in hypothalamic *Bdnf* and GABAergic markers gene expression, alongside decreased *Gfap*, suggesting a coordinated remodeling of glial activity, inhibitory tone and neural plasticity during chronic Semaglutide treatment. The source of this inhibitory input is still to be determined, but recent findings have shown that GABAergic TRH^Arc^ neurons directly suppress the activity of AgRP neurons, respond to GLP-1RA and contribute to its weight loss effects^13^. GABAergic *LepR/pDYN*^*+*^ neurons from the ventral dorsomedial nucleus of the hypothalamus are also selective afferents to AgRP neurons and suppress their activity under food availability^20^, suggesting that local hypothalamic circuits are the source of the altered inhibitory inputs described in the current work.

AgRP neurons are highly adapted to perceive and respond to negative energy balance, integrating hormonal, nutritional and sensory information to coordinate feeding behavior and metabolic homeostasis^6^. Our data shows that Semaglutide decreases energy expenditure (Fig. 1D), as well as O_2_ (Fig. 1F) and CO_2_ levels (Fig. 1G), indicative of a hypometabolic state. A potential reasoning for the paradoxical activation of AgRP neurons under sustained Semaglutide treatment is that these cells can perceive the drug-induced energy deficit, signaling the need to mobilize fat storages, which ultimately culminates in weight loss. Our data showing that blockade of adrenergic β3 receptors, expressed mainly in the adipose tissue and activated by the sympathetic nervous system^18,22^, has no effect in acute Semaglutide-induced weight loss, but inhibits the sustained one aligns with the idea of recruitment of lipolysis as a mechanism to restore homeostasis (Fig. S2). A complementary hypothesis is that hormonal responses evoked by semaglutide-induced negative energy balance, such as potential increase in ghrelin and decrease in leptin levels, might contribute to AgRP activation. However, we ruled out that hypothesis by showing that Semaglutide does not affect ghrelin plasma, and that leptin co-treatment with semaglutide enhances rather than inhibits weight loss (See Fig. S1D and 1F), suggesting that AgRP activation is not simply a compensatory response to hormonal alterations.

We have previously shown that hypothalamic energy circuits undergo dynamic synaptic remodeling in response to energy status and hormones through the involvement of neuron-glia interactions^23-26^, and that astrocytes in the hypothalamus express the GLP-1 receptor and regulate systemic glucose homeostasis^27^. Astrocytes are known to remodel their interactions with AgRP neurons in response to metabolic state^24, 25, 28-30^. Our data showing alterations in *Gfap* expression suggests that Semaglutide engages astrocyte-mediated mechanisms to remodel AgRP circuitry over time, thereby progressively shifting the dependence on AgRP neurons to sustain its metabolic effects, a hypothesis that warrants direct investigation.

Our findings also demonstrate that the requirement of AgRP neurons for Semaglutide-induced weight loss is modulated by dietary composition. While AgRP neurons are required for sustained weight loss in lean mice on a standard chow diet, this requirement is abolished in obese mice or in lean mice acutely exposed to HFD. Previous reports demonstrated that consumption of HFD bypass AgRP regulation of feeding and relies mainly on dopaminergic signaling^16,17^, suggesting that HFD shifts Semaglutide’s primary site of action from homeostatic hypothalamic AgRP circuits to hedonic feeding pathways, such as dopaminergic neurons in the VTA (17), the Lateral Hypothalamus^31^, or the Nucleus Tractus Solitarius^32,33^. Given that GLP-1R activation suppresses hedonic feeding^34,35^, Semaglutide may primarily engage hedonic pathways under HFD conditions rather than hypothalamic energy balance circuits. This may explain why some individuals on GLP-1 therapy report reduced cravings and altered food preferences rather than purely suppressed hunger^36,37^. A recent study has reported that ablation of *Glp1r*^+^ neurons in the arcuate nucleus does not impair Semaglutide-induced weight loss^14^, suggesting that this region might be dispensable for this effect. However, it is important to highlight that AgRP neurons do not express GLP-1 receptors^10,11^ and would not have been captured by that ablation model. Our findings add a complementary perspective by showing that the necessity of AgRP neurons for Semaglutide-induced weight loss is both time- and context-dependent, becoming more critical with prolonged treatment and under specific dietary conditions. Notably, the importance of *Glp1r*^+^ neurons in different brain areas, particularly those projecting from NTS to the paraventricular hypothalamus, was characterized in male mice maintained on high-fat diet^14^, a context in which we also observe no dependence on AgRP neurons for weight loss promoted by Semaglutide. Together, these findings support a model in which Semaglutide engages distinct neurocircuits depending on dietary and hormonal state, with a progressive recruitment of hypothalamic pathways, such as AgRP circuitry. This convergence highlights the importance of considering treatment duration, sex, and metabolic context when dissecting the neural mechanisms of GLP-1–based therapies.

A notable finding was that three days of HFD exposure was sufficient to eliminate AgRP neuron dependence, suggesting that even short-term dietary changes may alter GLP-1RA neural targets. This has clinical implications for individuals undergoing dietary transitions, such as those shifting from a Western diet to a lower-fat diet, or individuals cycling between different macronutrient compositions^38,39^. If diet composition influences how GLP-1R agonists act on different neural pathways, responses to therapy may vary based on habitual diet and recent dietary history. Moreover, switching mice from HFD back to a standard diet restored AgRP neuron dependence, indicating that the effects of dietary fat exposure on GLP-1R signaling are reversible. This suggests that dietary composition dynamically regulates energy balance circuits, potentially influencing long-term weight loss maintenance. Understanding whether similar transitions in humans affect GLP-1 therapy efficacy is an important avenue for future research.

The observed sex differences in AgRP neuron dependence suggest that hormonal and metabolic factors may differentially influence GLP-1R agonist efficacy in males and females. Sexual hormones are known to modulate hypothalamic circuits^40-43^, including AgRP neurons, and may enhance sensitivity to Semaglutide’s effects on energy balance in females. Further research is needed to determine whether sex hormones directly influence GLP-1R signaling in AgRP neurons or whether alternative pathways, such as differential engagement of reward circuits or peripheral metabolic adaptations, contribute to the observed differences.

A potential limitation of our model is that AgRP hypofunction alters the baseline biochemical and hormonal background of KO mice^44^, which could contribute to the loss of Semaglutide’s effect. However, our data showing that short-term HFD exposure alone is sufficient to alter Semaglutide’s mechanism of action suggests that diet, rather than baseline metabolic differences, is the primary driver of this shift. While it remains possible that AgRP-*Sirt1* KO mice have unique metabolic adaptations that influence their response to Semaglutide, the fact that dietary composition alone modifies its mechanism argues against metabolic background being the sole determinant.

Overall, our findings show for the first time that Semaglutide’s mechanism of action is not static but dynamically influenced by dietary composition. Under standard chow conditions, weight loss requires AgRP neurons and homeostatic energy regulation, while under HFD conditions, Semaglutide engages dopaminergic circuits to suppress hedonic feeding behavior, overriding the need for AgRP neuron involvement. These insights may have clinical implications, highlighting the potential for diet to alter GLP-1-based therapeutic strategies for obesity and metabolic disorders. We also challenge the current idea that Semaglutide inhibits AgRP activity, showing that sustained treatment leads to AgRP activation which is critical for sustaining its weight loss effects.

## Methods

### Mice

All animals (4-8 weeks old) were housed under standard laboratory conditions on a 12-hour light/12-hour dark cycle (Lights on at 7AM) with *ad libitum* access to food and water, and controlled ambient temperature. Animals were group housed for maintanence and single caged either in standard animal facility cages or in Promethion™ metabolic and behavioral phenotyping system during experimental procedures.

AgRP^DTR^ or wild-type mice from the same litter were injected with diphtheria toxin (50 µg/kg, subcutaneous) between P3-P5 and ablation was confirmed by immunohistochemistry for beta-galanin, as previously reported^9,37^. AgRP-*Sirt1* KO lines have been described elsewhere^36^, but in summary transgenic mice expressing *Cre* recombinase selectively in the *Agrp*-expressing cells were bred with mice harboring a targeted mutant *Sirt1* allele (Sirt1^*loxP*^, B6; 129-Sirt1^*tm1gu*^/J; stock number 008041 – The Jackson Laboratory) flanking the exon of the *Sirt1* gene, which encodes Sirt1 catalytic domain. When bred with the Tg.AgrpCre mice (B6.129S4-Gt(ROSA)26Sor^*tm1Sor*^— originally from The Jackson Laboratory), the deleted *Sirt1* allele (Sirt1^Δex4^) transcribes a mutant protein that has no apparent residual Sirt1 activity or dominant negative effects. AgRP^tdTomato^ animals were obtained by crossing Tg.AgrpCre with *tdTomato*^*(Ai14)*^-floxed animals purchased from Jackson’s Lab (Stock number 007914). C57BL/6J mice were purchased from Jackson’s Lab (Stock number 000664).

Mice had access to either Standard chow diet (Inotiv, 2018SC; % kcal: 18.6 protein, 6.2 fat, 44.2 carbohydrate, energy density of 3.1 kcal/g) or High-Fat Diet (Research Diets, D12492; % kcal: 20.1 protein, 19.6 carbohydrate, 60 fat, energy density of 5.21 kcal/g). For diet-induced obesity experiments, animals (6-9 weeks old at the start of the experiment) were single housed for at least 12 weeks with *ad libitum* access to High-Fat Diet and water. Body weight was measured weekly and animals that gained more than 30% of weight were considered obese, as previously published^25^. Body composition was measured using NMR technology (EchoMRI; Echo Medical Systems). Animal husbandry, housing, and euthanasia were performed according to the protocol reviewed and approved by the Yale University Institutional Animal Care and Use Committee and in compliance with the NIH Guide for the Care and Use of Laboratory Animals.

### Drugs and reagents

Mice were intraperitoneally injected with a drug or it’s correspondent vehicle during the Light Cycle (7 AM – 7 PM). Semaglutide Sodium Salt (Cayman Chemical, 40170, 160 ng/g or 40nmol/kg/day) was dissolved in PBS and administered in a dose like the ones reported previously in the literature^46^. SR 59230-A (Sigma-Aldrich, St. Louis, S8688, 0.5 mg/kg) was dissolved in PBS + DMSO 1.6%. Leptin (Thermo Fisher Scientific, 450-31-1MG, 1 mg/kg) was diluted in PBS.

### Whole-cell patch clamp recordings

Mice were anesthetized with isoflurane and decapitated around 10 AM, the brain was rapidly removed and immersed in a cold (4° C) and oxygenated cutting solution containing (mM): sucrose 220, KCl 2.5, NaH_2_PO_4_ 1.23, NaHCO_3_ 26, CaCl_2_ 1, MgCl_2_ 6 and glucose 10 (pH 7.3 with NaOH). Coronal hypothalamic slices (300 µm thick) were cut with a Leica vibratome after the brain was trimmed to a small tissue block containing the hypothalamus. After preparation, slices were maintained at room temperature (23–25° C) in a storage chamber in the artificial cerebrospinal fluid (ACSF) (bubbled with 5% CO_2_ and 95% O_2_) containing (in mM): NaCl 124, KCl 3, CaCl_2_ 2, MgCl_2_ 2, NaH_2_PO_4_ 1.23, NaHCO_3_ 26, glucose 10 (pH 7.4 with NaOH) for recovery and storage. After recovery at room temperature for at least 1 hour, slices were transferred to a recording chamber constantly perfused with bath solution (same as the ACSF except containing 2.5 mM glucose) at a temperature of 33° C and a perfusion rate of 2 ml/min for electrophysiological experiments. Whole-cell patch clamp recording was performed in *Agrp-tdTomato* neurons under current clamp, as previously reported^47^. Spontaneous and miniature excitatory and inhibitory currents in the same cell were recorded for 3-5 minutes for both vehicle- and semaglutide-treated mice. For evaluation of miniature currents, cells were incubated with tetrodotoxin (TTX, 1 µM) Frequency (Hz) of currents was analyzed using AxoGraph 4.9.

### Indirect calorimetry analysis

Mice were individually housed in metabolic cages (Promethion Core®, Sable Systems, North Las Vegas, USA) with temperature- and humidity-controlled system (maintained at ∼21°C and 40-50% relative humidity). Animals were acclimated to the metabolic chambers for at least two days prior to experimental manipulations. The Promethion system continuously measured oxygen consumption and carbon dioxide production. Airflow was maintained at 2.0 L/min, and reference air samples were collected at regular intervals to account for background fluctuations. Respiratory exchange ratio (RER) was calculated as the ratio of VCO_2_ to VO_**2**_. Energy expenditure (EE) was determined using the Weir equation. Food and water intake were monitored using gravimetric sensors, allowing for real-time quantification of consumption. Caloric intake was determined from food consumption data (Considering energy density of SD and HFD), and energy balance was calculated as the difference between energy intake and total energy expenditure. Real time data for body weight was collected through an in-cage enrichment device every time the animal interacts with it. Whole-body fat utilization was calculated using the follow equation: 1.67 × (VO_2_ − VCO_2_). Whole-body carbohydrate utilization was calculate using the follow equation: 4.55 × VCO_2_ – 3.21 × VO_2_, as described elsewhere^18^. Locomotor activity was continuously recorded using an XYZ beam-break system integrated into the calorimetry chambers, which measured total and ambulatory activity. Ambulatory activity is defined here as a goal-oriented behavior computed every time there is an increase in speed that lasts for more than 2 seconds. Raw data was processed through Sable Systems Macro Interpreter v.24.10.2 and data was analyzed using CalR 1.3 (48).

### Immunofluorescence

Brains were fixed through transcardiac perfusion with aldehyde fixative (4% paraformaldehyde + 15% picric acid in 0.1 M PB). Samples were left overnight in 4% paraformaldehyde. Brains were cut into 50 μm thick coronal sections using a Leica VT1000P vibratome throughout the whole caudal-rostral extent of the hypothalamus (20-30 sections per brain). Free-floating sections were permeabilized with Triton X-100 0.15% (v/v) for 15 minutes and incubated in blocking serum (1% Bovine serum albumin) for one hour at room temperature followed by an overnight incubation with primary rat anti-cfos (1:1000, #226 017, Synaptic Systems, Goettingen, Germany). Sections were washed three times with 0.1M PB and then incubated for 90 minutes with secondary antibody (Alexa Fluor 488-conjugated goat anti-guinea pig, 1:2000, Thermofisher #A11073). Sections were washed three times in 0.1M PB and then mounted on slides with Vectashield containing DAPI and then imaged and processed on a Zeiss Axioplan2 fluorescent scope (×20). Stack confocal images (6 μm, 1024 × 1024) were manually analysed (5 sections per mouse per experiment) to calculate the amount of co-localization between c-fos and td-Tomato signal.

### RNA isolation and quantification through RT-qPCR

Animals were deeply anesthetized with isoflurane and killed by decapitation. Hypothalamic tissue was collected and frozen in liquid nitrogen. Total RNA was extracted using RNeasy® lipid mini kit (Qiagen, 74106). NanoDrop 2000 spectrophotometer (Thermo Fisher Scientific) was used to quantify the amount of RNA in each sample and assessment of A260/A280 and A260/A230 ratios were used to determine potential contamination by protein or guanidine, respectively. cDNA was reverse transcribed (Qiagen, 205311) and amplified with SYBR Green Supermix (Bio-Rad) using a QuantStudio 6 Pro Real-Time PCR System (Applied Biosystems). Calculations were performed by a comparative method (2ΔΔCT) and relative expression was adjusted to *G3pdh* housekeeping gene. The following primers were used:

### Metabolic and endocrine assessment

Blood samples were collected from awake mice via the retro-orbital sinus using a heparinized capillary tube. Mice were manually restrained, and blood was drawn swiftly to minimize stress. Approximately 100-150 µL of blood was collected per mouse. Blood glucose and ketone bodies were measured through commercial detection strips (Precision Xtra). Following collection, blood samples were immediately transferred to microcentrifuge tubes and allowed to clot at room temperature for 30 minutes. Samples were then centrifuged at 10.000 × g for 10 minutes at 4°C to separate the serum. The supernatant (serum) was carefully transferred to a new tube and stored at -80°C until further analysis. Serum leptin, insulin and ghrelin levels were measured through a commercial ELISA kit (Crystal Chem, leptin: # 90030, insulin #90080, and ghrelin: Millipore #EZRGRA-90K) following the manufacturer’s instructions. Briefly, serum samples were thawed on ice, diluted as needed, and loaded into pre-coated ELISA plates. Standards and controls were prepared according to kit protocols. Absorbance was measured at the specified wavelength using a microplate reader, and sample concentrations were interpolated from the standard curve.

### Statistical analysis

All statistical analysis were run in GraphPad Prism 10.2.3. Comparisons between two groups were done through the t-test. Analysis of variance (ANOVA) was used to compare multiple groups. When assessing the interaction between genotype (AgRP-Sirt1 Control vs KO) and treatment (Vehicle vs Semaglutide), data was analyzed through two-way ANOVA followed by Šidák or Tukey *post-hoc*. Data normality was assessed through D’Agostino-Pearson omnibus and Shapiro-Wilkin tests. Details on the statistical analyses can be found at the Supplementary Table 1, containing sample size and further information on statistical analysis of each experiment in which we found statistical difference. p<0.05 was considered statistically significant.

## Ethics declaration – Competing interests

The authors declare no competing interests.

## Data availability

All data needed to interpret, verify and extend the research discussed in the article are available in the main text, supplementary materials or the source data. Any additional data can be provided under request to the corresponding author.

## Acknowledgments

We thank M. Shanabrough for technical assistance to develop the current work. MD thank A. M. Herculano, K. R. H. M. Oliveira, and F. Borba for the discussions and training that led to this study.

## Funding

CAPES-Fulbright Full PhD Fellowship, grant CAPES call 05/2021 number 88881.625374/2021-01 (MD).

## Author contributions

Conceptualization: MD, TLH, Methodology: MD, RCP, TLH, Investigation: MD, RCP, ZWL, Visualization: MD, Supervision: TLH, Writing—original draft: MD, Writing—review & editing: MD, TLH

## Notes

### Competing Interest Statement

The authors have declared no competing interest.

