## Supplementary material for "Diet context gates AgRP neuron involvement in Semaglutide-induced weight loss": BioRXiv - Supplementary Information.pdf

Mateus d'Ávila *et al.*

**This PDF file includes:**

Supplementary Text
Figs. S1 to S2
Tables S1

**Fig. S1.**

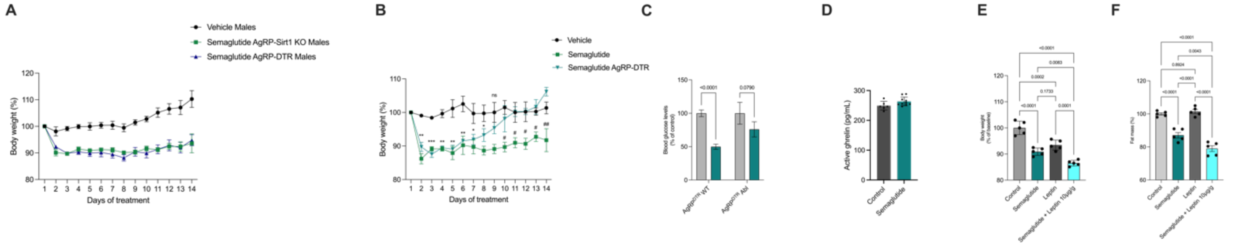


**AgRP neurons are dispensable in male lean mice and hormonal alterations do not play a role in Semaglutide-induced weight loss. (A)** Male AgRP-*Sirt1* KO or AgRP$^{DTR}$ mice treated with Semaglutide show no weight rebound across 14 days of injections, **(B)** Female AgRP$^{DTR}$ show weight rebound after 8 days of daily injections (n = 10), **(C)** Blood glucose levels of AgRP$^{DTR}$ wild-type (Controls) and ablated (Abl) showing that absence of AgRP neurons impair hypoglycemic effects of Semaglutide (n = 6; Two-way ANOVA followed by Šidák *post-hoc*, p-values showed in figure), **(D)** Ghrelin levels in the plasma of control vehicle vs semaglutide treated mice (Control n = 6, Semaglutide = 10; Welch's t-test, p-value above 0.05), **(E)** Body weight of female mice treated with PBS (Control, n = 5), Semaglutide (n = 5), Leptin (10 µg/g, n = 5) or Semaglutide+Leptin (n = 5). One-way ANOVA followed by *Tukey* post-hoc, p-values displayed in the figure, **(F)** Levels of fat mass in PBS (Control, n = 5), Semaglutide (n = 5), Leptin (10 µg/g, n = 5) or Semaglutide+Leptin (n = 5). One-way ANOVA followed by *Tukey* post-hoc, p-values displayed in the figure. ***$p<0.001$, **$p<0.01$, *$p<0.5$ vs Vehicle.

**Fig. S2.**

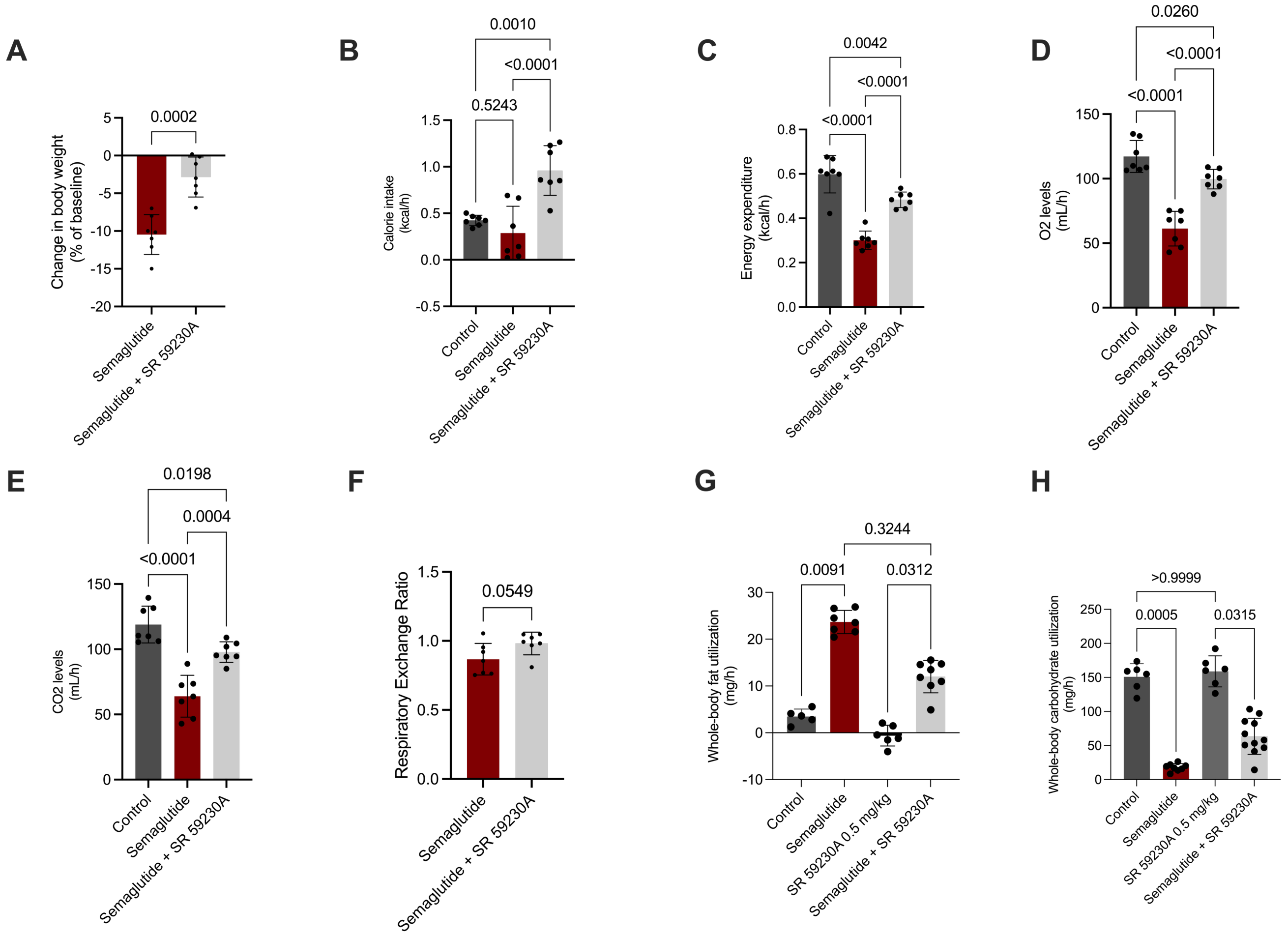


**Blockade of beta3 adrenergic receptor prevents Semaglutide-induced weight loss effects and metabolic actions.** **(A)** Percent change in body weight in animals pre-treated with the β3-adrenergic receptor antagonist SR 59230A and treated with Semaglutide, **(B)** Calorie intake (kcal/h) of animals pre-treated with the β3-adrenergic receptor antagonist SR 59230A and treated with Semaglutide, **(C)** Energy expenditure (kcal/h) of animals pre-treated with the β3-adrenergic receptor antagonist SR 59230A and treated with Semaglutide,**(D)** $O_2$ and **(E)** $CO_2$ levels of animals pre-treated with the β3-adrenergic receptor antagonist SR 59230A and treated with Semaglutide, **(F)** Respiratory exchange ratio of animals pre-treated with the β3-adrenergic receptor antagonist SR 59230A and treated with Semaglutide. Silver bars represent the period in which mice received Semaglutide treatment, **(G)** Whole-body fat utilization, and **(H)** Whole-body carbohydrate utilization. Details on statistical analysis can be found at supplementary Table

**Fig. S3.**

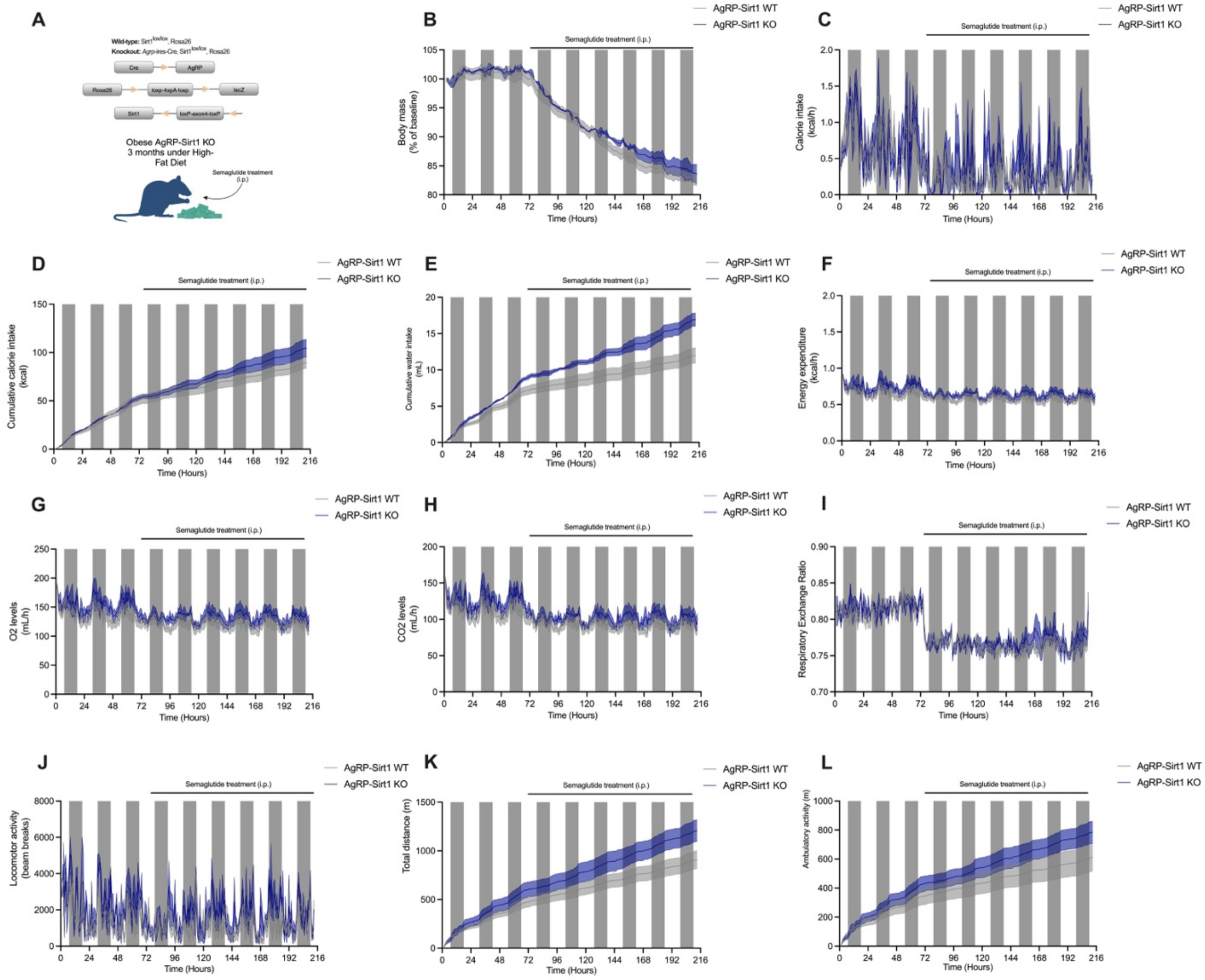


**AgRP neurons are dispensable in obese male mice. (A)** Schematic representation of AgRP-*Sirt1* Control Wild-type (WT, n = 5) and Knockout (KO, n = 6) obese mice under High-Fat Diet for 3 months before start of Semaglutide injections, **(B)** Body mass (% of baseline) of animals treated with Semaglutide, **(C)** Calorie intake (kcal/h) of animals treated with Semaglutide, **(D)** Cumulative food intake (kcal) of animals, **(E)** Cumulative water intake of WT and KO mice, **(F)** Energy expenditure (kcal/h) of animals before and during Semaglutide treatment, **(G)** $O_2$ levels of animals before and during Semaglutide treatment, **(H)** $CO_2$ levels of animals treated with Semaglutide, **(I)** Respiratory exchange ratio of animals before and during Semaglutide treatment, **(J)** Locomotor activity (beam breaks) of WT and KO animals before and during Semaglutide treatment, **(K)** Total distance (m) of WT and KO animals across days of treatment, **(L)** Ambulatory activity (m) of WT and KO animals across days of treatment.

**Table S1. Summary of statistical analysis performed in the study.**

| Figure | Sample size (n) | Statistical test | Statistical Test summary (p-value,F, df, t, and/or ANOVA table) | Notes |
|---|---|---|---|---|
| 1B | AgRP-Sirt1 CTL and KO = 15 | Welch's t-test, two-tailed | p=0.0197; t=2.499; df=23.93 | % of weight loss after 2 days of Semaglutide treatment. |
| 1B | AgRP-Sirt1 CTL and KO = 15 | Welch's t-test, two-tailed | p=<0.0001; t=11.94; df=22.56 | % of weight loss after 15 days of Semaglutide treatment. |
| 1B | AgRP-Sirt1 CTL = 6<br>KO = 7 | Two-way ANOVA, Šidák *post-hoc* | ANOVA table SS (Type III) DF MS F (DFn, DFd) P value<br>Interaction 0.3626 1 0.3626 F (1, 22) = 1.822 P=0.1908<br>Treatment 23.81 1 23.81 F (1, 22) = 119.6 P<0.0001<br>Genotype 18.38 1 18.38 F (1, 22) = 92.31 P<0.0001<br>Residual 4.380 22 0.1991 | Fat mass levels of AgRP-Sirt1 animals after 2 days of injection with Semaglutide |
| 1B | AgRP-Sirt1 CTL = 6<br>KO = 7 | Two-way ANOVA, Šidák *post-hoc* | ANOVA table SS (Type III) DF MS F (DFn, DFd) P value<br>Interaction 22.83 1 22.83 F (1, 22) = 34.58 P<0.0001<br>Treatment 4.526 1 4.526 F (1, 22) = 6.854 P=0.0157<br>Genotype 0.02382 1 0.02382 F (1, 22) = 0.03607 P=0.8511<br>Residual 14.53 22 0.6603 | Fat mass levels of AgRP-Sirt1 animals after 15 days of injection with Semaglutide |
| 1C | AgRP-Sirt1 CTL and KO = 10 | Two-way ANOVA, Šidák *post-hoc* | ANOVA table SS DF MS F (DFn, DFd) P value<br>Interaction 1520 1 1520 F (1, 36) = 9.490 P=0.0039<br>Treatment 128421 1 128421 F (1, 36) = 801.6 P<0.0001<br>Genotype 3095 1 3095 F (1, 36) = 19.32 P<0.0001<br>Residual 5767 36 160.2 | Cumulative calorie intake (kcal) of AgRP-Sirt1 CTL and KO mice in baseline vs treatment. |
| 1I | AgRP-Sirt1 CTL and KO = 8 | Two-way ANOVA, Tukey *post-hoc* | ANOVA table SS DF MS F (DFn, DFd) P value<br>Interaction 3212 1 3212 F (1, 28) = 36.72 P<0.0001<br>Genotype 41.86 1 41.86 F (1, 28) = 0.4785 P=0.4948<br>Treatment 93.16 1 93.16 F (1, 28) = 1.065 P=0.3109<br>Residual 2449 28 87.48 | Glucose levels of AgRP-Sirt1 Control vs Knockout mice treated with vehicle or Semaglutide for 15 days. AgRP-Sirt1 Control Vehicle vs AgRP-Sirt1 KO Semaglutide p-value = 0.99. |
| 1J | AgRP-Sirt1 CTL and KO = 8 | Two-way ANOVA, Tukey *post-hoc* | ANOVA table SS DF MS F (DFn, DFd) P value<br>Interaction 0.04147 1 0.04147 F (1, 28) = 11.65 P=0.0020<br>Genotype 0.06813 1 0.06813 F (1, 28) = 19.14 P=0.0002<br>Treatment 0.09743 1 0.09743 F (1, 28) = 27.37 P<0.0001<br>Residual 0.09969 28 0.003560 | Insulin levels of AgRP-Sirt1 Control vs Knockout mice treated with vehicle or Semaglutide for 15 days. |
| 1K | AgRP-Sirt1 CTL Vehicle = 5<br>AgRP-Sirt1 CTL Semaglutide = 6<br>AgRP-Sirt1 KO Vehicle = 5<br>AgRP-Sirt1 KO Semaglutide = 6 | Two-way ANOVA, Tukey *post-hoc* | ANOVA table SS (Type III) DF MS F (DFn, DFd) P value<br>Interaction 143.0 1 143.0 F (1, 18) = 278.9 P<0.0001<br>Genotype 9.703 1 9.703 F (1, 18) = 18.92 P=0.0004<br>Treatment 0.4794 1 0.4794 F (1, 18) = 0.9349 P=0.3464<br>Residual 9.231 18 0.5128 | Leptin levels of AgRP-Sirt1 Control vs Knockout mice treated with vehicle or Semaglutide for 15 days. |
| 2A | AgRP-Sirt1 CTL Vehicle 2 days = 3<br>AgRP-Sirt1 CTL Semaglutide 2 days = 4<br>AgRP-Sirt1 CTL Vehicle 15 days = 3<br>AgRP-Sirt1 CTL Semaglutide 15 days = 5 | Two-way ANOVA, Tukey *post-hoc* | ANOVA table SS (Type III) DF MS F (DFn, DFd) P value<br>Interaction 79683 1 79683 F (1, 11) = 15.37 P=0.0024<br>Time of treatment 77656 1 77656 F (1, 11) = 14.98 P=0.0026<br>Treatment 76175 1 76175 F (1, 11) = 14.70 P=0.0028<br>Residual 57021 11 5184 | *Agrp* gene expression of animals treated with Semaglutide for 2 or 15 days (dpi). |
| 2A | AgRP-Sirt1 CTL Vehicle 2 days = 3<br>AgRP-Sirt1 CTL Semaglutide 2 days = 4<br>AgRP-Sirt1 CTL Vehicle 15 days = 3<br>AgRP-Sirt1 CTL Semaglutide 15 days = 5 | Two-way ANOVA, Tukey *post-hoc* | ANOVA table SS (Type III) DF MS F (DFn, DFd) P value<br>Interaction 42855 1 42855 F (1, 11) = 24.04 P=0.0005<br>Time of treatment 27215 1 27215 F (1, 11) = 15.27 P=0.0024<br>Treatment 11210 1 11210 F (1, 11) = 6.288 P=0.0291 | *Npy* gene expression of animals treated with Semaglutide for 2 or 15 days (dpi). |

<table>
<tr><td></td><td></td><td></td><td>Residual 19611 11 1783</td><td></td></tr>
<tr><td>2B</td><td>Control and Semaglutide = 5</td><td>Unpaired t-test, two-tailed</td><td>t=4.690, df=8</td><td>c-fos/AgRP colocalization in slices of animals treated with Semaglutide for 15 days.</td></tr>
<tr><td>2C</td><td>Control = 10<br>Semaglutide = 13</td><td>Welch’s t-test, two-tailed</td><td>t=2.848; df=18</td><td>sIPSC of AgRP neurons from animals treated for 2 days.</td></tr>
<tr><td>2C</td><td>Control = 9<br>Semaglutide = 10</td><td>Welch’s t-test, two-tailed</td><td>t=3.409, df=16</td><td>mIPSC of AgRP neurons from animals treated for 2 days.</td></tr>
<tr><td>2C</td><td>Control = 10<br>Semaglutide = 11</td><td>Welch’s t-test, two-tailed</td><td>t=4.148, df=12</td><td>sEPSC of AgRP neurons from animals treated for 2 days.</td></tr>
<tr><td>2C</td><td>Control = 8<br>Semaglutide = 8</td><td>Welch’s t-test, two-tailed</td><td>t=0.445, df=14</td><td>mEPSC of AgRP neurons from animals treated for 2 days.</td></tr>
<tr><td>2D</td><td>Control = 10<br>Semaglutide = 11</td><td>Welch’s t-test, two-tailed</td><td>t=2.316, df=18</td><td>sIPSC of AgRP neurons from animals treated for 15 days.</td></tr>
<tr><td>2D</td><td>Control = 10<br>Semaglutide = 11</td><td>Welch’s t-test, two-tailed</td><td>t=1.28, df=18</td><td>mIPSC of AgRP neurons from animals treated for 15 days.</td></tr>
<tr><td>2D</td><td>Control = 10<br>Semaglutide = 10</td><td>Welch’s t-test, two-tailed</td><td>t=0.7242, df=12</td><td>sEPSC of AgRP neurons from animals treated for 15 days.</td></tr>
<tr><td>2D</td><td>Control = 9<br>Semaglutide = 11</td><td>Welch’s t-test, two-tailed</td><td>t=0.458, df=16.7</td><td>mEPSC of AgRP neurons from animals treated for 15 days.</td></tr>
<tr><td>2E</td><td>WT Vehicle: 4<br>WT Semaglutide: 5<br>KO Vehicle: 5<br>KO Semaglutide: 6</td><td>Two-way ANOVA, Sidak post-hoc</td><td>
<table>
<tr><th>Šídák's multiple comparisons test</th><th>Predicted (LS) mean diff.</th><th>95.00% CI of diff.</th><th>Below threshold?</th><th>Summary</th><th>Adjusted P Value</th></tr>
<tr><td>slc17a6</td><td></td><td></td><td></td><td></td><td></td></tr>
<tr><td>WT Vehicle vs. WT Semaglutide</td><td>-0.1050</td><td>-1.312 to 1.102</td><td>No</td><td>ns</td><td>>0.9999</td></tr>
<tr><td>WT Vehicle vs. KO Vehicle</td><td>0.5759</td><td>-0.6311 to 1.783</td><td>No</td><td>ns</td><td>0.7455</td></tr>
<tr><td>WT Vehicle vs. KO Semaglutide</td><td>0.5167</td><td>-0.6447 to 1.678</td><td>No</td><td>ns</td><td>0.8010</td></tr>
<tr><td>WT Semaglutide vs. KO Vehicle</td><td>0.6809</td><td>-0.4571 to 1.819</td><td>No</td><td>ns</td><td>0.5092</td></tr>
<tr><td>WT Semaglutide vs. KO Semaglutide</td><td>0.6218</td><td>-0.4678 to 1.711</td><td>No</td><td>ns</td><td>0.5641</td></tr>
<tr><td>KO Vehicle vs. KO Semaglutide</td><td>-0.05917</td><td>-1.149 to 1.030</td><td>No</td><td>ns</td><td>>0.9999</td></tr>
<tr><td>gabra1</td><td></td><td></td><td></td><td></td><td></td></tr>
<tr><td>WT Vehicle vs. WT Semaglutide</td><td>-0.7949</td><td>-2.002 to 0.4121</td><td>No</td><td>ns</td><td>0.3956</td></tr>
<tr><td>WT Vehicle vs. KO Vehicle</td><td>-0.7152</td><td>-1.922 to 0.4919</td><td>No</td><td>ns</td><td>0.5207</td></tr>
<tr><td>WT Vehicle vs. KO Semaglutide</td><td>-0.4398</td><td>-1.601 to 0.7217</td><td>No</td><td>ns</td><td>0.8946</td></tr>
<tr><td>WT Semaglutide vs. KO Vehicle</td><td>0.07972</td><td>-1.058 to 1.218</td><td>No</td><td>ns</td><td>>0.9999</td></tr>
<tr><td>WT Semaglutide vs. KO Semaglutide</td><td>0.3551</td><td>-0.7344 to 1.445</td><td>No</td><td>ns</td><td>0.9457</td></tr>
<tr><td>KO Vehicle vs. KO Semaglutide</td><td>0.2754</td><td>-0.8142 to 1.365</td><td>No</td><td>ns</td><td>0.9843</td></tr>
<tr><td>gabra3</td><td></td><td></td><td></td><td></td><td></td></tr>
<tr><td>WT Vehicle vs. WT Semaglutide</td><td>-0.6685</td><td>-1.876 to 0.5386</td><td>No</td><td>ns</td><td>0.5977</td></tr>
<tr><td>WT Vehicle vs. KO Vehicle</td><td>0.06827</td><td>-1.139 to 1.275</td><td>No</td><td>ns</td><td>>0.9999</td></tr>
<tr><td>WT Vehicle vs. KO Semaglutide</td><td>-0.4226</td><td>-1.584 to 0.7389</td><td>No</td><td>ns</td><td>0.9111</td></tr>
<tr><td>WT Semaglutide vs. KO Vehicle</td><td>0.7367</td><td>-0.4013 to 1.875</td><td>No</td><td>ns</td><td>0.4157</td></tr>
<tr><td>WT Semaglutide vs. KO Semaglutide</td><td>0.2459</td><td>-0.8436 to 1.335</td><td>No</td><td>ns</td><td>0.9913</td></tr>
<tr><td>KO Vehicle vs. KO Semaglutide</td><td>-0.4908</td><td>-1.580 to 0.5987</td><td>No</td><td>ns</td><td>0.7918</td></tr>
<tr><td>slc32a1</td><td></td><td></td><td></td><td></td><td></td></tr>
<tr><td>WT Vehicle vs. WT Semaglutide</td><td>-2.777</td><td>-3.984 to -1.570</td><td>Yes</td><td>****</td><td><0.0001</td></tr>
<tr><td>WT Vehicle vs. KO Vehicle</td><td>0.3710</td><td>-0.8360 to 1.578</td><td>No</td><td>ns</td><td>0.9587</td></tr>
<tr><td>WT Vehicle vs. KO Semaglutide</td><td>-0.9889</td><td>-2.150 to 0.1726</td><td>No</td><td>ns</td><td>0.1382</td></tr>
<tr><td>WT Semaglutide vs. KO Vehicle</td><td>3.148</td><td>2.010 to 4.286</td><td>Yes</td><td>****</td><td><0.0001</td></tr>
<tr><td>WT Semaglutide vs. KO Semaglutide</td><td>1.788</td><td>0.6983 to 2.877</td><td>Yes</td><td>***</td><td>0.0002</td></tr>
<tr><td>KO Vehicle vs. KO Semaglutide</td><td>-1.360</td><td>-2.449 to -0.2704</td><td>Yes</td><td>**</td><td>0.0067</td></tr>
<tr><td>bdnf</td><td></td><td></td><td></td><td></td><td></td></tr>
<tr><td>WT Vehicle vs. WT Semaglutide</td><td>-1.231</td><td>-2.438 to -0.02402</td><td>Yes</td><td>*</td><td>0.0431</td></tr>
<tr><td>WT Vehicle vs. KO Vehicle</td><td>0.2341</td><td>-0.9730 to 1.441</td><td>No</td><td>ns</td><td>0.9962</td></tr>
<tr><td>WT Vehicle vs. KO Semaglutide</td><td>-0.1135</td><td>-1.275 to 1.048</td><td>No</td><td>ns</td><td>>0.9999</td></tr>
<tr><td>WT Semaglutide vs. KO Vehicle</td><td>1.465</td><td>0.3271 to 2.603</td><td>Yes</td><td>**</td><td>0.0048</td></tr>
<tr><td>WT Semaglutide vs. KO Semaglutide</td><td>1.118</td><td>0.02799 to 2.207</td><td>Yes</td><td>*</td><td>0.0413</td></tr>
<tr><td>KO Vehicle vs. KO Semaglutide</td><td>-0.3476</td><td>-1.437 to 0.7420</td><td>No</td><td>ns</td><td>0.9508</td></tr>
<tr><td>gfap</td><td></td><td></td><td></td><td></td><td></td></tr>
<tr><td>Vehicle - Semaglutide</td><td></td><td></td><td></td><td></td><td></td></tr>
<tr><td>AgRP-Sirt1 WT</td><td>0.7120</td><td>0.3865 to 1.038</td><td>Yes</td><td>***</td><td>0.0001</td></tr>
<tr><td>AgRP-Sirt1 KO</td><td>0.06968</td><td>-0.2242 to 0.3635</td><td>No</td><td>ns</td><td>0.8124</td></tr>
<tr><td>Vehicle - Semaglutide</td><td></td><td></td><td></td><td></td><td></td></tr>
<tr><td>AgRP-Sirt1 WT</td><td>0.7120</td><td>0.3865 to 1.038</td><td>Yes</td><td>***</td><td>0.0001</td></tr>
<tr><td>AgRP-Sirt1 KO</td><td>0.06968</td><td>-0.2242 to 0.3635</td><td>No</td><td>ns</td><td>0.8124</td></tr>
<tr><td>aif1</td><td></td><td></td><td></td><td></td><td></td></tr>
<tr><td>WT Vehicle vs. WT Semaglutide</td><td>0.2434</td><td>-0.9636 to 1.450</td><td>No</td><td>ns</td><td>0.9953</td></tr>
<tr><td>WT Vehicle vs. KO Vehicle</td><td>0.6355</td><td>-0.5715 to 1.843</td><td>No</td><td>ns</td><td>0.6519</td></tr>
<tr><td>WT Vehicle vs. KO Semaglutide</td><td>0.6621</td><td>-0.4994 to 1.824</td><td>No</td><td>ns</td><td>0.5653</td></tr>
<tr><td>WT Semaglutide vs. KO Vehicle</td><td>0.3921</td><td>-0.7459 to 1.530</td><td>No</td><td>ns</td><td>0.9301</td></tr>
<tr><td>WT Semaglutide vs. KO Semaglutide</td><td>0.4187</td><td>-0.6709 to 1.508</td><td>No</td><td>ns</td><td>0.8879</td></tr>
<tr><td>KO Vehicle vs. KO Semaglutide</td><td>0.02659</td><td>-1.063 to 1.116</td><td>No</td><td>ns</td><td>>0.9999</td></tr>
</table>
</td><td></td></tr>
<tr><td>S2A</td><td>Semaglutide = 6<br>Semaglutide + SR 59230A = 6</td><td>Welch’s t-test, two-tailed</td><td>t=5.386, df=12</td><td>Change in body weight of animals pre-treated with adrenergic beta3 antagonist SR 69230A</td></tr>
<tr><td>S2B-2F</td><td>Control = 7<br>Semaglutide = 7<br>Semaglutide+SR 59230A = 7</td><td>One-way ANOVA, Tukey post-hoc</td><td></td><td>Metabolic assessment of animals treated with Semaglutide for 7 days and then treated with the antagonist+Semaglutide. P-values displayed in the figures.</td></tr>
</table>

<table>
<tr><td>S2G</td><td>Vehicle: 5<br>Semaglutide: 7<br>SR-59230A: 6<br>Semaglutide+SR59<br>230A: 11</td><td>One-way ANOVA,<br>Kruskal-Wallis post-hoc</td><td><table>
<tr><th>Dunn's multiple comparisons test</th><th>Mean rank diff.</th><th>Significant?</th><th>Summary</th><th>Adjusted P Value</th></tr>
<tr><td>Control vs. Semaglutide</td><td>-14.20</td><td>Yes</td><td>**</td><td>0.0091</td></tr>
<tr><td>Control vs. SR 59230A 0.5 mg/kg</td><td>4.967</td><td>No</td><td>ns</td><td>>0.9999</td></tr>
<tr><td>Control vs. Semaglutide + SR 59230A</td><td>-6.575</td><td>No</td><td>ns</td><td>0.7895</td></tr>
<tr><td>Semaglutide vs. SR 59230A 0.5 mg/kg</td><td>19.17</td><td>Yes</td><td>****</td><td><0.0001</td></tr>
<tr><td>Semaglutide vs. Semaglutide + SR 59230A</td><td>7.625</td><td>No</td><td>ns</td><td>0.3244</td></tr>
<tr><td>SR 59230A 0.5 mg/kg vs. Semaglutide + SR 59230A</td><td>-11.54</td><td>Yes</td><td>*</td><td>0.0312</td></tr>
</table></td><td></td></tr>
<tr><td>S2H</td><td>Vehicle: 5<br>Semaglutide: 7<br>SR-59230A: 6<br>Semaglutide+SR59<br>230A: 11</td><td>One-way ANOVA,<br>Kruskal-Wallis post-hoc</td><td><table>
<tr><th>Dunn's multiple comparisons test</th><th>Mean rank diff.</th><th>Significant?</th><th>Summary</th><th>Adjusted P Value</th></tr>
<tr><td>Control vs. Semaglutide</td><td>19.42</td><td>Yes</td><td>***</td><td>0.0005</td></tr>
<tr><td>Control vs. SR 59230A 0.5 mg/kg</td><td>-1.667</td><td>No</td><td>ns</td><td>>0.9999</td></tr>
<tr><td>Control vs. Semaglutide + SR 59230A</td><td>11.21</td><td>No</td><td>ns</td><td>0.0906</td></tr>
<tr><td>Semaglutide vs. SR 59230A 0.5 mg/kg</td><td>-21.08</td><td>Yes</td><td>***</td><td>0.0001</td></tr>
<tr><td>Semaglutide vs. Semaglutide + SR 59230A</td><td>-8.205</td><td>No</td><td>ns</td><td>0.3128</td></tr>
<tr><td>SR 59230A 0.5 mg/kg vs. Semaglutide + SR 59230A</td><td>12.88</td><td>Yes</td><td>*</td><td>0.0315</td></tr>
</table></td><td></td></tr>
<tr><td>3C</td><td>AgRP-Sirt1 Control<br>Vehicle = 8<br>AgRP-Sirt1 Control<br>Semaglutide = 8<br>AgRP-Sirt1 KO<br>Vehicle = 8<br>AgRP-Sirt1 KO<br>Semaglutide = 9</td><td>Two-way ANOVA,<br>Tukey post-hoc</td><td><table>
<tr><th>ANOVA table</th><th>SS (Type III)</th><th>DF</th><th>MS</th><th>F (DFn, DFd)</th><th>P value</th></tr>
<tr><td>Interaction</td><td>0.1092</td><td>1</td><td>0.1092</td><td>F (1, 29) = 0.005340</td><td>P=0.9422</td></tr>
<tr><td>Treatment</td><td>1794</td><td>1</td><td>1794</td><td>F (1, 29) = 87.77</td><td>P<0.0001</td></tr>
<tr><td>Genotype</td><td>427.8</td><td>1</td><td>427.8</td><td>F (1, 29) = 20.93</td><td>P<0.0001</td></tr>
<tr><td>Residual</td><td>592.9</td><td>29</td><td>20.44</td><td></td><td></td></tr>
</table></td><td>Fat mass of obese female mice treated with Semaglutide for 7 days.</td></tr>
<tr><td>4B</td><td>AgRP-Sirt1 CTL and KO= 11</td><td>Unpaired t-test, two-tailed</td><td>p=<0.0001, t=6.16; df= 20</td><td>Data comparing endpoint of both groups.</td></tr>
<tr><td>4H</td><td>AgRP-Sirt1 CTL and KO= 11</td><td>Unpaired t-test, two-tailed</td><td>p=0.0004; t=4.21;df=20</td><td>Data comparing 48 hours after Semaglutide injection.</td></tr>
</table>
